# LBFextract: unveiling transcription factor dynamics from liquid biopsy data

**DOI:** 10.1101/2024.05.03.592314

**Authors:** Isaac Lazzeri, Benjamin Gernot Spiegl, Samantha O. Hasenleithner, Michael R. Speicher, Martin Kircher

**Affiliations:** Institute of Human Genetics, Diagnostic and Research Center for Molecular BioMedicine, Medical University of Graz, Neue Stiftingtalstr. 6, 8010, Graz, Austria; Division of Oncology, Department of Internal Medicine, Medical University of Graz, 8010 Graz, Austria; BioTechMed-Graz, Graz, Austria; Berlin Institute of Health (BIH) at Charite – Universita tsmedizin Berlin, 10178, Berlin, Germany; Institute of Human Genetics, University Medical Center Schleswig-Holstein, University of Lu beck, Lu beck, Germany

## Abstract

**Motivation:** The analysis of circulating cell-free DNA (cfDNA) holds immense promise as a non-invasive diagnostic tool across various human conditions. However, extracting biological insights from cfDNA fragments entails navigating complex and diverse bioinformatics methods, encompassing not only DNA sequence variation but also epigenetic characteristics like nucleosome footprints, fragment length, and methylation patterns.

**Results:** We introduce LBFextract, a comprehensive package designed to streamline feature extraction from cfDNA data, with the aim of enhancing the reproducibility and comparability of liquid biopsy studies. LBFextract facilitates the integration of preprocessing and postprocessing steps through alignment fragment tags and a hook mechanism. It incorporates various methods, including coverage-based and fragment length-based approaches, alongside two novel feature extraction methods: an entropy-based method to infer TF activity from fragmentomics data and a technique to amplify signals from nucleosome dyads. Additionally, it implements a method to extract condition-specific differentially active TFs based on these features for biomarker discovery. We demonstrate the use of LBFextract for the subtype classification of advanced prostate cancer patients using coverage signals at transcription factor binding sites from cfDNA. We show that LBFextract can generate robust and interpretable features that can discriminate between different clinical groups. LBFextract is a versatile and user-friendly package that can facilitate the analysis and interpretation of liquid biopsy data.

**Data and Code Availability and Implementation:** LBFextract is freely accessible at https://github.com/Isy89/LBF. It is implemented in Python and compatible with Linux and Mac operating systems. Code and data to reproduce these analyses have been uploaded to 10.5281/zenodo.10964406.

**Contact:** For further information, contact.

**Supplementary Information:** For additional details see Supplementary Information. For usage of the package, refer to https://lbf.readthedocs.io/.

## IINTRODUCTION

Analyses of circulating cell-free DNA (cfDNA), i.e. the analysis of naturally occurring short DNA fragments in bodily fluids like blood and urine, are increasingly being adopted for the identification, diagnostic assessment and surveillance of various pathological and physiological states in humans (1–5). This gain in traction can be attributed to many factors that are fueling the growth of the cfDNA field, such as the increasing prevalence of cancer (6), rising preference for non-invasive procedures, various advantages of liquid biopsies, i.e. diagnostic approaches using samples of bodily fluids, over standard tissue biopsies, favorable government initiatives, and growing public and private interest. However, the bioinformatics approaches to harvesting the inherent biological information from cfDNA fragments are becoming more sophisticated and complex. Especially, as the cfDNA field begins to extend beyond the analysis of observed DNA sequence variation, such as single nucleotide variants (SNVs) and somatic copy number alterations (SCNAs). For example, several studies employing nucleosome position mapping have now provided evidence that cfDNA reflects nucleosome footprints and coverage profiles at regulatory regions such as transcription start sites (TSS) and transcription factor binding sites (TFBS) have been employed for the inference of gene expression, cancer detection and tissue deconvolution (7–10). Various research shows that the cellular nucleosomal architecture significantly influences DNA fragmentation, creating distinct patterns in not only the length of the fragments but also the frequency and type of specific motifs, which have also been used for cancer detection and classification (11–17). The extraction of epigenetic alterations, such as cfDNA fragment length, diverse cfDNA fragment patterns, methylation markers, and signals at open chromatin regions, will play an important role in the development of more advanced liquid biopsy technologies (18,19). Although this development of cfDNA-derived features is rapidly evolving and individual methods of feature extraction involve similar steps, a package that offers a straightforward and extendable collection of feature extraction methods, with the aim of enhancing experiment reproducibility and comparability, is currently lacking. While a variety of bioinformatics tools are available to calculate diverse types of genomic coverage measures, spanning from bin wise genomic read coverage like BEDtools (40,41) to region specific fragment coverage like DeepTools (36), in liquid biopsy, new methods aimed at enhancing the signal derived from nucleosome dyads have been proposed (9,14,20–22). However, these specialized methods are mostly provided as part of workflows, intertwined with preprocessing and GC bias correction steps, which hinders their reusability. In LBFextract we provide diverse feature extraction methods based on coverage. Further, we provide an easy way to integrate GC bias correction methods in form of alignment fragment tags, compatible with softwares like GCparagon (23), but tool agnostic and making this step uncoupled from the LBFextract feature extraction methods. To this end, in LBFextract, we implement commonly used coverage strategies like midpoint coverage, used in the Griffin package (21), and read coverage, proposed in (9), but generalizing them with a user-defined number of bases to retain from each fragment. We further implement a new method, which we call “coverage around dyads”, with the aim of enhancing the signal derived from nucleosome dyads through a better modelling of nucleosome position on DNA fragments. We also included several feature extraction methods to extract different types of fragment length distributions (FLDs) and fragment length ratios, which provide orthogonal information to coverage-based features as well as new entropy-based fragmentomics features. Through a hook mechanism, we provide entry points to integrate extra pre- and postprocessing steps, i.e., plugins, allowing to use third party software to customize the way reads are collected, how they are transformed, the process of signal extraction as well as the way genomic ranges are normalized. Finally, LBFextract provides a way to identify condition-specific statistically significant transcription factor signals and their enrichment analysis, enabling condition-specific biomarker discovery from liquid biopsy data. In the first part of this article, we present the package structure, followed by a description of the features extraction methods. In the final part, we provide a demonstration of a clinical use case of this package for the extraction of coverage signals at TFBSs for subtype classification in the context of advanced prostate cancer (Figure 1).

**Figure 1:**
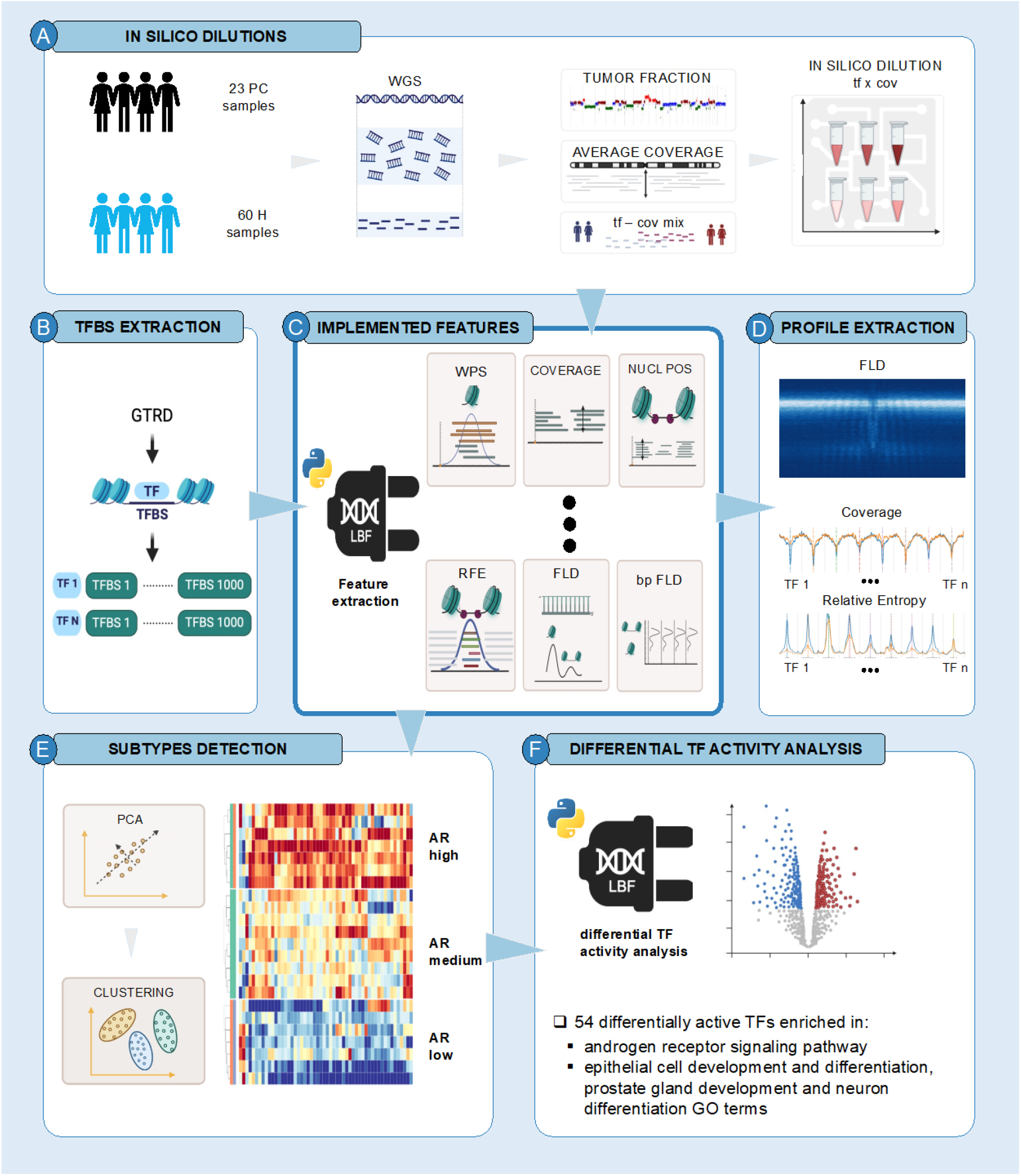
Workflow showing the application of LBFextract to PC subtype biomarker discovery. A) Prostate cancer samples are diluted to 20x coverage and 20% tumor fraction (Supplementary Info: *In silico* dilutions). B) TFBSs per TF are retrieved by the GTRD database. C) Coverage- and fragmentomics-based features are extracted from liquid biopsy data using LBFextract. D) Example of features extracted with LBFextract. E) Predefined set of features per TF are used and down projected to a lower dimensional space using PCA. This is followed by clustering to extract PC subtypes. F) Differential TF binding activity analyses are performed to extract condition-specific TFs. Further enrichment analyses are performed to place the detected differentially active TFs into context.

## MATERIALS AND METHODS

### Package Structure

LBFextract has been developed as a plugin system in which several hooks define entry points that a user can use to customize the workflow without having to reimplement code or functionality that the package or other plugins already implement. To achieve this, it uses Pluggy (24), a python package that allows the user to change the behavior of the host program. In this context, the modification of the behavior is defined as python functions called hooks, which are loaded and registered at runtime and change or exchange certain parts of the host program. We developed two types of hooks: Command Line Interface (CLI)-hooks and workflow-specific hooks. Using the CLI hooks, a user can implement CLI-plugin commands that are registered at installation time and are made available through the command line interface as LBFextract subcommands. The workflow-specific hooks allow the customization of different steps of the default workflow. Specifically, we implemented the following hooks: fetch_reads, save_reads, load_reads, transform_reads, transform_single_intervals, transform_all_intervals, save_signal, plot_signal and save_plot. The read fetching hook handles Binary Sequence Alignment Map (BAM) files, generally done by retrieving specific regions of interest defined in one or multiple Browser Extensible Data (BED) files. The way reads are saved and loaded is defined by the save_reads and the load_reads hooks. Signal extraction methods are implemented as transform_single_intervals hooks, which handles the extraction of the signal in each region defined in the BED file. The transform_all_intervals hook is available for transformations requiring all genomic intervals as input. The save_signal hook defines the way extracted signals are stored. Finally, signal-specific plots can be defined in the plot_signal hook and saved with the save_plot hook.

### Feature extraction methods

LBFextract defines a set of feature extraction methods, which can be divided into coverage and fragmentomics-based methods. Recent work has described diverse types of coverage signals that can be derived from cfDNA. Here, we implemented coverage (fragment coverage, midpoint coverage, middle-n points coverage, coverage around the dyad, sliding coverage, central 60bp-coverage) as well as fragmentomics signals (Windowed Protection Score (WPS), Fragment Length Distribution (FLD) and Fragment Length Ratios (FLR)). A central aspect of LBFextract is the introduction of a novel feature extraction method defined as Coverage Around Dyads and novel entropy-derived features such as Entropy and Relative Fragment Entropy (RFE).

To better describe the feature extraction methods, we introduce a mathematical notation for a BED file, DNA fragments and DNA fragments relative to a genomic interval.

We define a BED file as a multiset of genomic intervals *g*_*i*_:

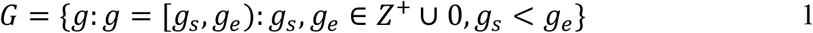

where *g*_*s*,_ *g*_*e*_ represent the start and end of a genomic interval *g*.

and the set of all fragments present in a sample as *F*:

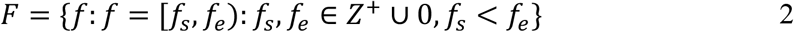

where *f*_*s*_ and *f*_*e*_ represent the start and end of fragments respectively. Further, let *g* be a target region, we define 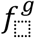 as the set of all positions of fragment *f* overlapping interval *g* expressed relative to interval *g*, which can be defined as:

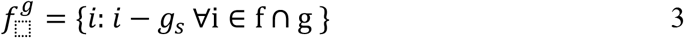

and 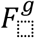 as the multiset of all 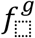 overlapping an interval *g*

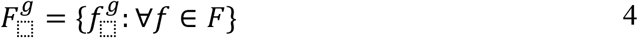

Hereafter, all intervals *g* will be considered to have the same length defined as *w*, which is by default set to 4000, which corresponds to ± 2000 bp around the TFBS center.

### Coverage

In fragment coverage, information concerning paired reads is used to infer the length of a fragment and the coverage at a specific position is defined as the number of fragments overlapping a given position. Here, positions covered by a fragment are those spanning from the left-most start to the right-most end of two mates in a read pair. This is different from read coverage, which considers positions of sequenced nucleotides, without taking overlaps or missing segments between the paired-end reads into account, thus possibly introducing an artifact. For any 0 ≤ *l* ≤ *w*, define 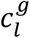 to be the coverage at position *l* in genomic interval *g* relative to interval *g* defined as:

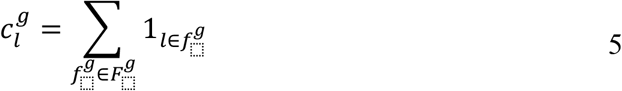

and ***c***^*g*^ ∈ *R*^*w*^ to be the coverage of interval g defined as:

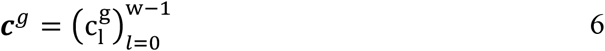

the coverage profile for all regions in a BED file can be described as:

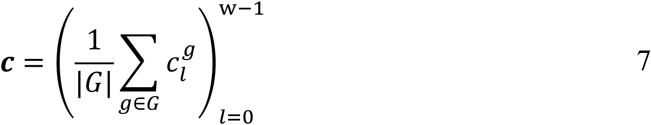

in which ***c*** ∈ *R*^*w*^.

Midpoint coverage is calculated as the number of fragments having their central position located at a certain genomic position. If 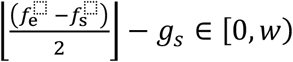 then define the midpoint to be:

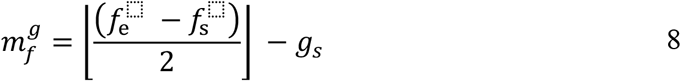

Then, the formula 5 changes to:

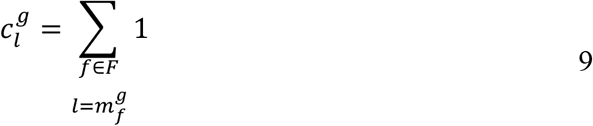

This measure might be problematic as coverage at each position is dramatically decreased by what could be considered an extreme *in silico* trimming of reads. Instead of midpoint coverage, *n* positions around the middle of each fragment might be used, which in LBFextract is called middle-n points coverage. To define this, let n be the number of positions around *m*, we can then describe 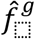 as the subset of *f* representing the positions around the midpoint of *f* and 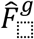 the multiset of all 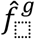 mapping to a genomic interval *g*:

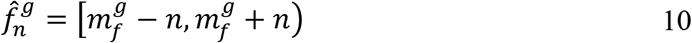

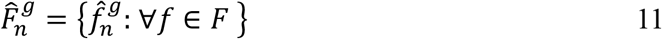

then the middle-n points coverage at position *l* with respect to genomic interval *g* can be defined as:

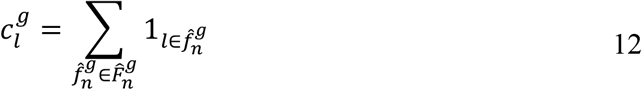

Midpoint coverage and middle-n points coverage have been widely used to extract information concerning nucleosome positioning (21). The middle points are assumed to be the location of the nucleosome dyads, which represent the most protected part of the DNA wrapped around nucleosomes. While this might generally be true for fragments smaller than 250 base pairs (bp), it is not true for longer fragments, i.e. >250bp. Indeed, the fragment midpoint of poly-nucleosomal structures like di-nucleosomes falls within the unprotected region between two nucleosomes interfering with proper nucleosome dyad localization. To avoid this problem, we implemented “coverage around dyads”. This method takes into consideration the presence of poly-nucleosomal structures and models the probability of each read coming from a n or n+1 poly-nucleosomal structure (Supplementary Information Algorithm 1 lines 7-8). This is used to reconstruct the size of each fragment before degradation (Supplementary Information Algorithm 1 lines 13-17), which in turn is used to better place the position of the dyad and obtain a stronger nucleosome derived signal (Supplementary Information Algorithm 1 lines 18-22). Fragment size of mono-nucleosomal-derived cfDNA is extracted from the fragment length distribution of the region chr12:34300000-34500000, which was described to show highly phased nucleosomes (Supplementary Information Algorithm 1 lines 1-2)(25).

Sliding coverage can be useful when the average depth of coverage is low to smooth the signal and reduce the effect of artifacts, such as drop in coverage at an individual position. The calculation uses a moving average over genomic intervals to calculate the coverage at each position:

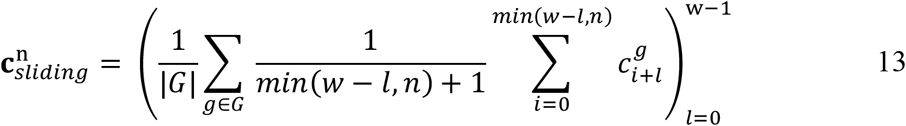

where *n* is the window parameter with a default of 4bp.

Central 60bp-coverage introduced in (9), trims 53 bases from both fragment sides and uses 60bp from each side for coverage calculation. We generalized this, introducing two variables (default [53, 113)) describing the range of bases that should be retained.

Profiles of each interval defined in the BED files are further normalized to the mean coverage of the flanking regions. Let *r* be the length of the flanking region, then the normalization step can be defined as follows:

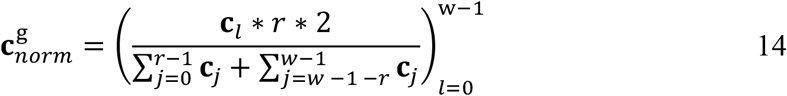

### Windowed Protection Score

Further, we implemented the Windowed Protection Score (WPS), introduced in (7), which quantifies the protective effect of nucleosomes in a genomic region by assigning a score to each position. This score is determined by the count of fragments that entirely cover the span of a window centered on a genomic position, subtracted by the number of fragments that begin/terminate within that window.

This can be represented as follow:

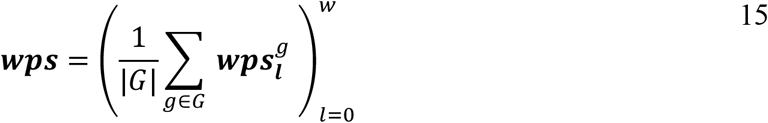

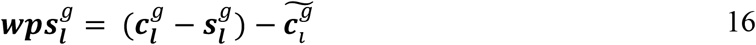

In which ***wps*** ∈ *R*^*w*^, ***c*** ∈ *R*^*w*^ is the coverage vector for genomic interval *g* calculated using a multiset of trimmed fragments corresponding to: {*f*: [*f*_*s*_ + *v, f*_*e*_ − *v*) ∀ *f* ∈ *F*}, *w* is the length of the genomic interval by default set to 4000, 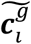 is the running median coverage vector, and ***s***^*g*^ defined as:

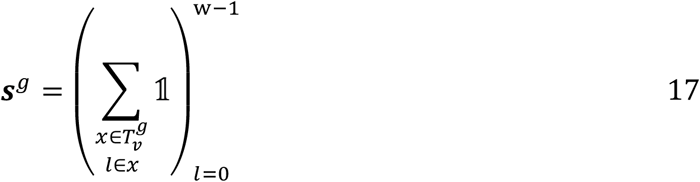

is the vector with the coverage of the genomic intervals corresponding to a window *v* surrounding the start and the end of each fragment, which are represented by the multiset *T*:

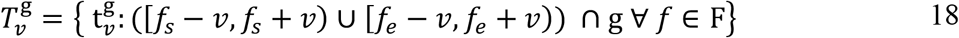

Profiles of each interval defined in the BED files are further normalized as done for the coverage-based signals (Formula 14).

### Fragment length distribution

In recent years, there has been a growing utilization of fragmentomics features. For example, the ratio between fragments having long and short sizes has been used for cancer prediction or for determining the proportion of placental cfDNA. To improve upon these methods, we provide the possibility to extract the full distribution of fragment lengths per position given the genomic intervals in one or multiple BED files. To calculate this, for all fragments *F* in BED file *G*, the fragment length distribution at position *i* ***d***_*i*_ can be defined as:

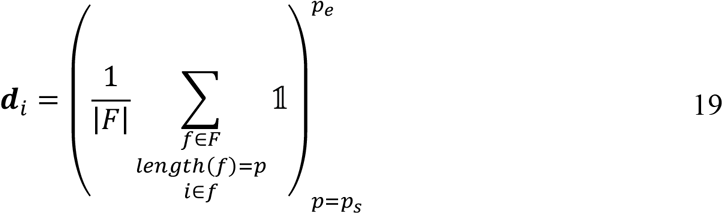

where *p*_*s*_ is the minimum fragment length to be considered and *p*_*e*_ the maximum. For each BED file containing the regions of interest, a matrix *D* = [***d***_0_, ***d***_1_, …, ***d***_*w*−1_] in which *w* represents the length of the regions, is generated. By description of fragments as a set of positions, we can define different types of fragment length distributions. For example, in “coverage around dyads” we described how inferred dyad locations are used in the set of positions rather than the fragment itself. The same principle can be applied to fragment length distribution feature extraction methods. In this way, we implemented the fragment length distribution around dyads, the fragment length distribution around the midpoint, the middle-n points fragment length distribution, and the central 60bp-fragment length distribution.

### Entropy and Relative Fragment Entropy

Previously, it was shown that fragmentation patterns at active transcription start sites (TSS) change, resulting in higher diversity of fragment lengths with respect to DNA protected regions. Prior work described a peak in the fragment length distribution around 160 bp as well as a correlation with RNA expression levels of individual genes (26). For this approach, termed epigenetic expression inference from cfDNA-sequencing (EPIC-seq), coverages of about 500x to 2000x are required, which are reached through hybrid capture-based targeted deep sequencing.

We hypothesized that higher diversity of fragment lengths may not only apply to TSSs, but also to TFBS, where nucleosome displacement and depletion occurs. Therefore, we defined a way to extract an entropy signal from genomic shotgun data as a measure of transcription factor activity. To overcome the problem of low depth of coverage at each position for typical sequencing data, we use multiple regions, such as multiple TFBSs representing the same TF. At a coverage of 20x, this results in an average number of ten thousand reads per position for TFs having 1,000 TFBSs along the genome. We implemented an entropy and a normalized entropy signal.

The former calculates the entropy of the fragment lengths at each position. Given a random variable *E* which takes values defined in the set *O* = {*i*: *i* ∈ [*p*_*s*_, *p*_*e*_)} distributed according to *P*_*l*_: *O* → ***d***_*l*_, entropy at position *l* can be defined as:

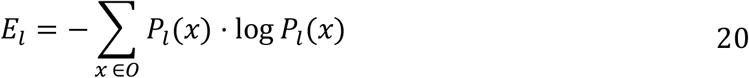

The latter, which we called Relative Fragment Entropy (RFE), calculates the divergence in the fragment length distribution at each position over genomic intervals and the fragment length distribution in the flanking regions.

This can be defined as:

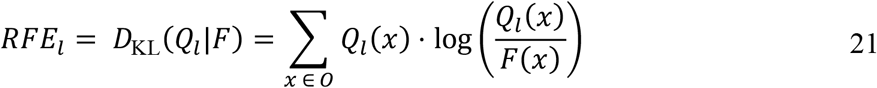

Where *l* represents the genomic position, *Q*_*l*_ represents the fragment length distribution at position *l* and *F* is the fragment length distribution in the flanking region.

At genuine TFBSs, we expect higher fragmentation than in the flanking regions where DNA is expected to be protected by nucleosomes. The FLD in the central part of the TFBS should differ from the FLD in the flanking region and therefore the RFE should show a peak in the center for TFs possessing higher activity (Supp. Figure 3).

To avoid effects due to differences in coverage between positions, the same number of reads is used to calculate the entropy and the RFE signals at each position.

### Differentially active genomic intervals

In LBFextract, we implemented a way to calculate differentially active genomic regions for which, in the case of activity, a peak or a dip is expected in the central part of the region. The general procedure is summarized in Algorithm 2 (Supplementary Information). In the first part, the algorithm extracts the features f for each transcription factor t from a sample’s BAM file (line 7-9). Subsequently, it calculates the accessibility of each feature and applies an appropriate statistical test (line 10-23). It uses the accessibility values of each TF grouped according to a label vector l, which assigns each sample to a specific group. Correction for multiple testing is applied to adjust for the higher probability of observing statistically significant results when testing multiple groups and multiple TFs. Finally, for all TFs, which were found to be differentially active, an enrichment analysis is retrieved through the string API.

## RESULTS

### Difference in androgen receptor signaling in prostate cancer and castration resistant prostate cancer

Our previous work utilized tissue and cancer type-specific chromatin accessibility datasets to identify tissue-dependent TFBS accessibility patterns and we found evidence that nucleosome footprints in cfDNA are informative of TFBSs (9). In addition, we demonstrated that TFs are amenable to molecular prostate cancer subtyping, which is an important issue in the management of prostate cancer (27). More specifically, we focused on the phenomenon of transdifferentiation of prostate adenocarcinoma (PRAD) to a treatment-emergent small-cell neuroendocrine prostate cancer (t-SCNC), which is a frequent mechanism in the development of treatment resistance against androgen deprivation therapy (ADT), and constitutes a subtype that is no longer dependent on androgen receptor (AR) signaling (28). The ability to detect this critical transition in longitudinal sampling has clinical implications, i.e. as it constitutes a change in therapy (27). The involvement of TFs in this transdifferentiation process to neuroendocrine prostate cancer (NEPC) has been extensively studied (28–30). Also, we have previously leveraged this information to confirm transdifferentiation events in our prostate cancer cohort (9). Herein, we use LBFextract to reproduce our previous finding and extend them by extracting not only coverage but also fragmentomic features at diverse TFBSs of TFs involved in this trans-differentiation. We apply the differential activity analysis provided by LBFextract to shed light on subtype-specific TFs. To demonstrate the validity of these signals and their potential, we showcase a previously described patient P148 (8). Within 12 months, the time interval between collection of the samples P148_1 and P148_3, the prostate adenocarcinoma transdifferentiated to a t-SCNC, which was accompanied by a clinical observation of a decrease of prostate-specific antigen (PSA) and an increase of neuron-specific enolase (NSE). We look at AR chromatin accessibility via coverage and FLDs at TFBSs of AR for P148. To reduce confounding effects like tumor fraction and coverage, which may bias the analysis, we additionally performed in silico dilutions for both samples to 20x coverage and 20% tumor fraction (Supplementary Info: *In silico* dilutions). We retrieved TFBSs from the Gene Transcription Regulation Database (GTRD v21.12) sorted them based on the number of peaks supporting each meta-cluster, removed TFBSs belonging to sex-related chromosomes and retained the top 1,000 TFBSs. In doing so, we obtained 1058 TFs with 1,000 TFBSs each (Figure 1).

Through the analysis of general coverage at AR-specific TFBSs conducted with the LBFextract extract_coverage method, we observed the expected central increase in the normalized coverage in P148_3 relative to P148_1, showing the reduced AR activity (Figure 2a-e). By analyzing the coverage signal with the extract_coverage_around_dyads, we were also able to observe the peaks flanking the central positions of the TFBSs in the case of P148_1, which indicates the presence of nucleosome phasing, which was not observed using normal fragment coverage. Furthermore, these peaks were found to be absent for P148_3, suggesting reduced AR binding to the TFBSs and therefore a decreased phasing of the neighboring nucleosomes (Figure 2b).

**Figure 2:**
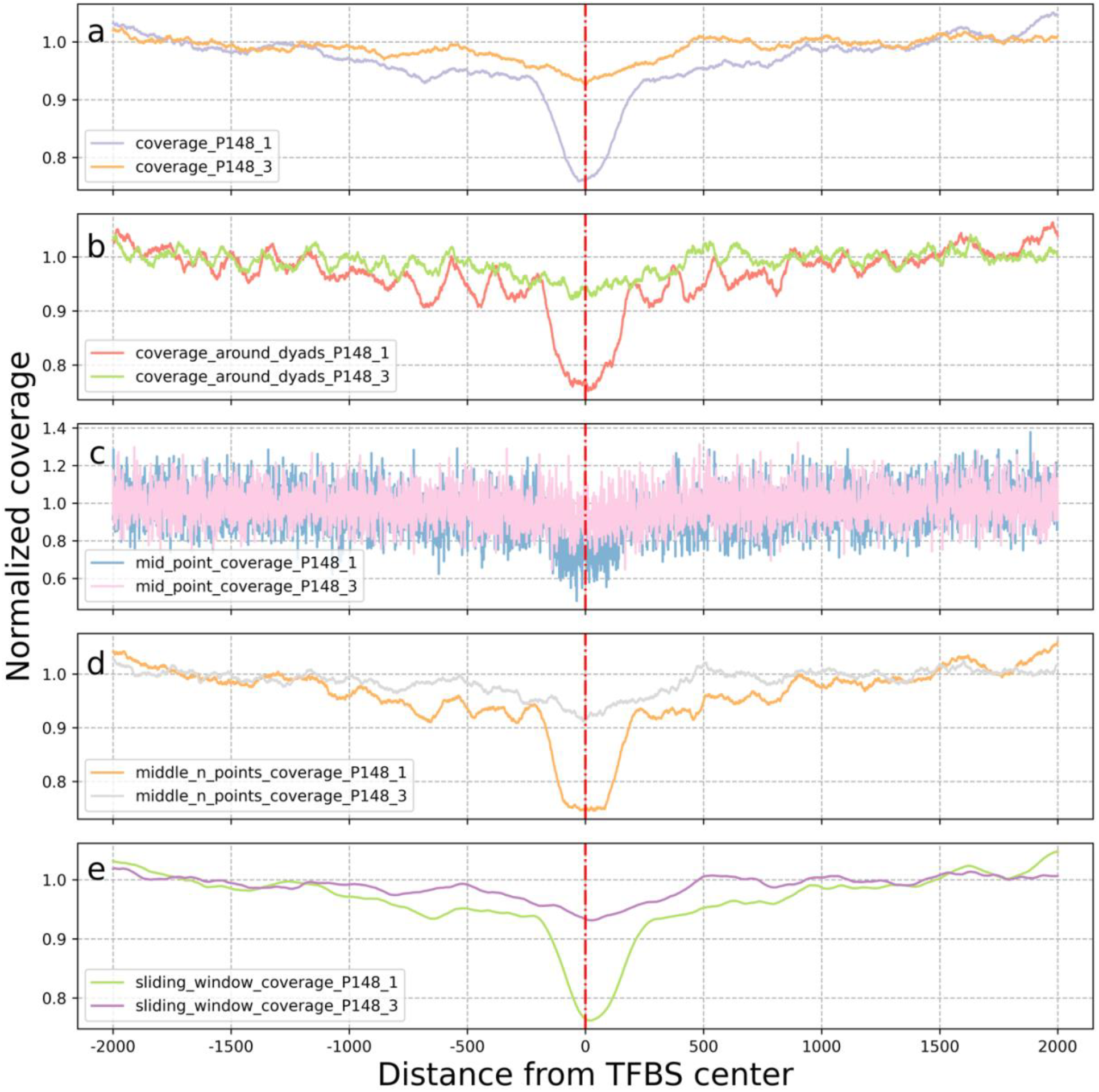
Comparison of coverage signals of cfDNA samples obtained for the same individual at two time points (P148_1 and P148_3). a) Normal coverage b) Coverage around dyads c) Midpoint coverage d) Middle-n points coverage e) Sliding window coverage. The figure shows the respective overlays of 1000 sites for the AR TF and highlights a difference between these samples around the central location of the considered windows, suggesting regions of open chromatin in P148_1 that are absent in P148_3.

The same could be observed from fragmentomics features. When looking at the FLD 2000bp around the TFBS of AR (Figure 3), from which we subtracted the signal of the FLD in the flanking regions (fFLD), an increased diversity in fragmentation patterns towards the center of the TFBS at fragment lengths above 180 bp can be observed in P148_1 (Figure 3a), which is almost absent in P148_3 (Figure 3b). Furthermore, a decreased representation of the fragment lengths around 166 bp toward the center of the TFBS is visible in the case of P148_1 (light blue blob in Figure 3a), while less pronounced in P148_3. Similarly, this information is captured by relative entropy (Figure 4, Figure 3c), which exploits the difference between the distribution in the flanking regions and the one at each position. A higher relative entropy value at AR for sample P148_1 is evidence for a higher diversity in fragment lengths at the center of the TFBSs. Because peaks or valleys are not dependent on coverage here, but only on the fragment length, the information contained in the fragment length distribution or entropy-derived signals offers a unique perspective on TF-specific TFBSs’ chromatin state.

**Figure 3:**
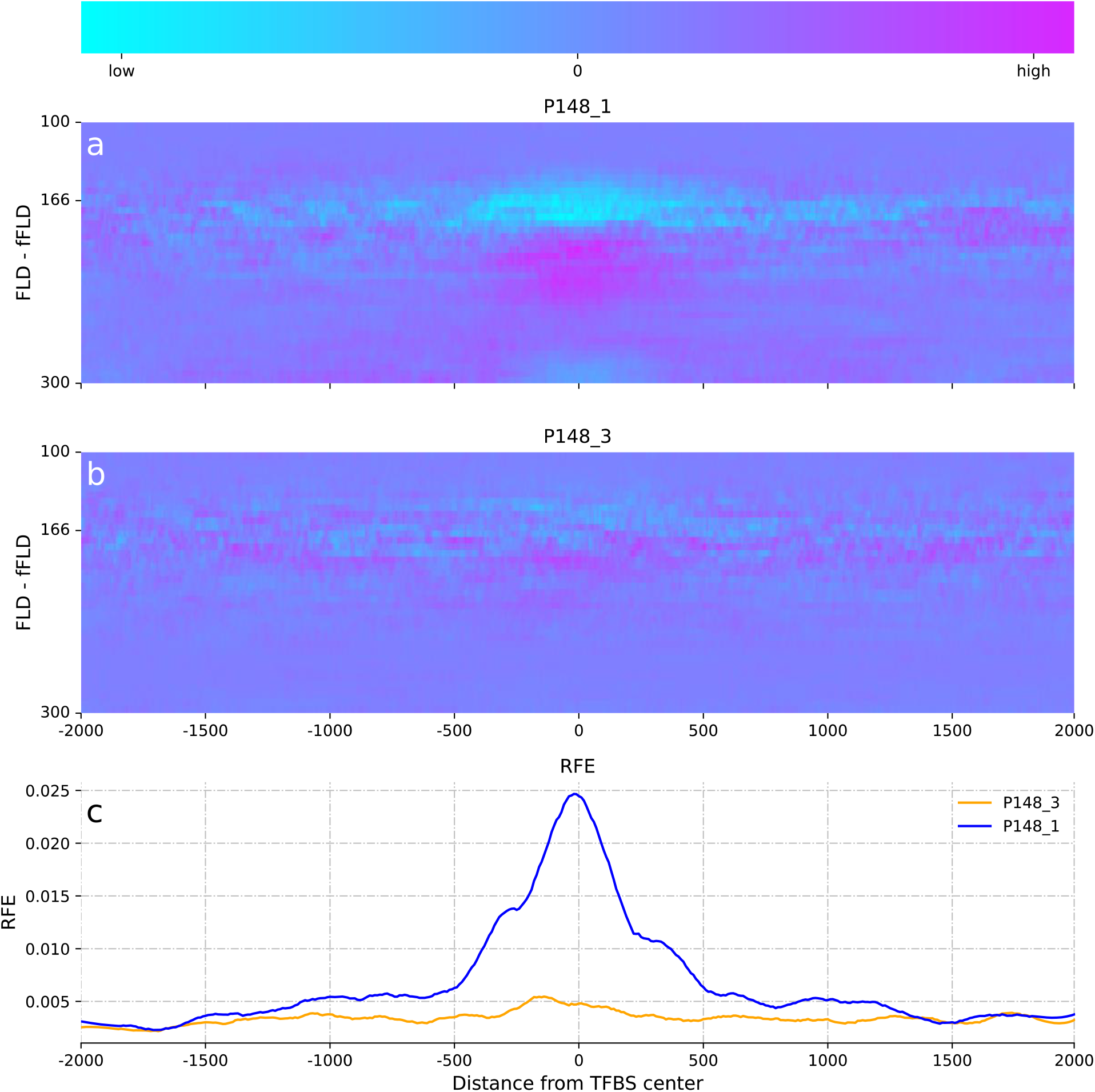
Fragment length distributions (FLD) per position minus the FLD in the flanking regions (fFLD) at androgen receptor (AR)-specific TFBSs. a) FLD - fFLD P148_1. b) FLD - fFLD P148_3. c) RFE per position at AR-specific TFBSs smoothed with Savitzky-Golay filter.

**Figure 4:**
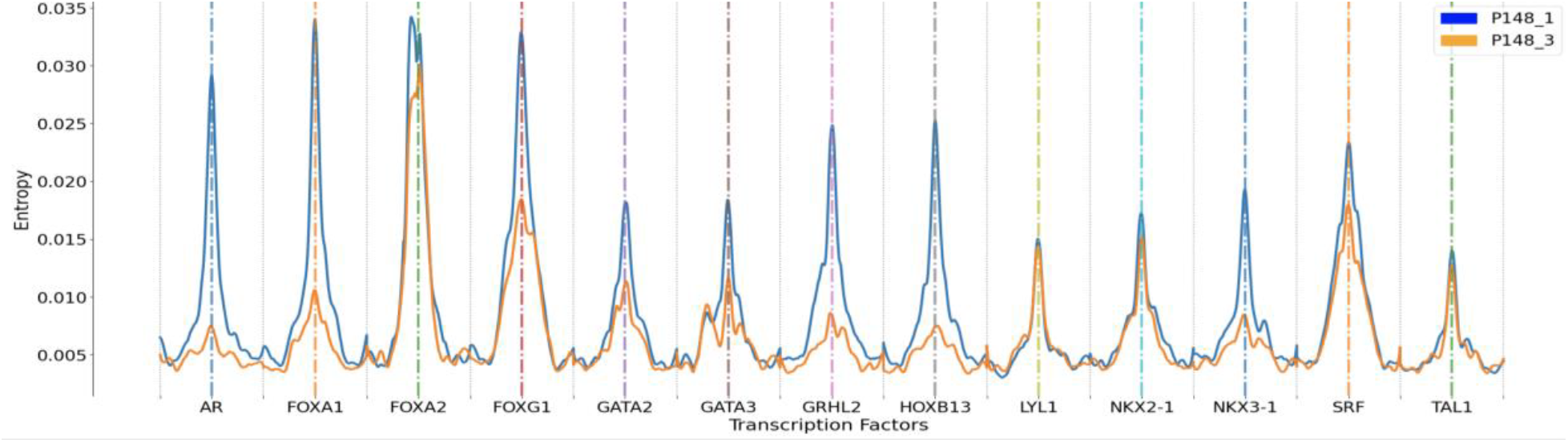
Relative Fragment Entropy for TFs involved in the transdifferentiation process. (i.e., AR, FOXA1/2, FOXG1, GATA2/3, GRHL2, HOXB13, NKX2-1, NKX3-1, SRF) as well as TFs involved in hematopoiesis (LYL1, TAL1).

**Figure 5:**
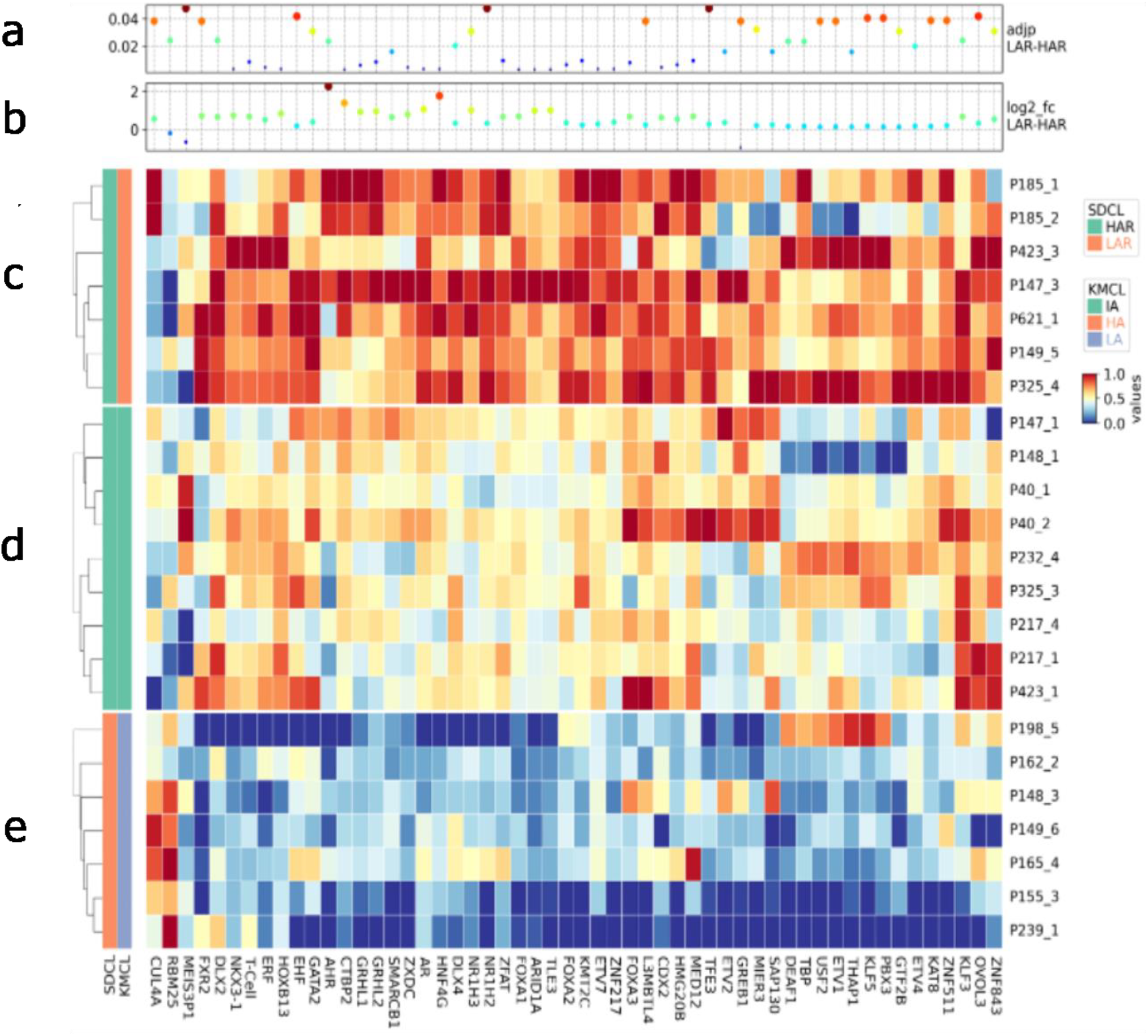
Heatmap of the accessibilities of differentially active TFs found in the HAR and LAR clusters. a) Dot-plot of the adjusted p-values. b) Dot-plot of the pseudo log2 fold changes. c-e) Heat maps of the signals in the HAR (c, d) and LAR clusters (e). Bars on the left represent the clustering according to semisupervised clustering (SDCL) and kmeans (KMCL), respectively.

### Characterization of androgen receptor high and low signal-specific TFs in advanced prostate cancer

Despite the presence of samples with high and low AR signals, it is essential to emphasize that the signal patterns are not uniform and sampled from at least two distributions. Further, we postulate the existence of prostate cancer subtypes characterized by high and low AR signals within our prostate cancer cohort. To further explore this theory, we applied semi-supervised and k-means clustering methods to detect different subgroups finding 2 and 3 clusters respectively. With this aim, in the semi-supervised approach, we selected samples P148_1 (PRAD condition) and P148_3 (NEPC condition), which display high and low AR coverage signals, respectively, as prototypes for High AR (HAR) and Low AR (LAR) clusters. To make use of prior knowledge, we expanded the sets of TFs with those found to be differentially active in NEPC and PRAD in (20) and (8). We obtained a final set of TFs for the neuroendocrine condition composed of: AR, FOXA1, NKX3-1, HOXB13, GRHL2, ASCL1, GATA2 and HNF4G. Each prostate cancer sample was assigned to its nearest cluster prototype (lowest Euclidean distance), giving rise to an HAR cluster (n=15), characterized by the presence of a valley around the TFBSs centers of AR (Supp. Figure 5a), FOXA1 (Supp. Figure 5b), NKX3-1 (Supp. Figure 5c), HOXB13 (Supp. Figure 5d), GRHL2 (Supp. Figure 5f) and HNF4G (Supp. Figure 5i) and a LAR cluster (n = 7) characterized by flatter profiles for the same TFs. We extracted coverage signals for all 1058 TFs and assessed differences in TF activities between clusters using the Mann-Whitney U test. After correction for multiple testing using the Benjamini-Hochberg method, we rejected all null hypotheses with an adjusted-p value lower than 0.05. From the first analysis between HAR and LAR groups, we found 53 TFs with significantly increased accessibility (Table 1). Most of the TFs nominated for the initial clustering dataset—with the exception of ASCL1—were also found to be present within the list of differentially active TFs, which supports the validity of the initial set of TFs. Concurrently, we observe genes which were previously shown to be linked to the AR CRPC subtype, such as AR and FOXA1, at the top of the list of differentially active TFs. AR, which has a statistically relevant 1.06 log2 fold downregulation in the LAR cluster, belongs to the steroid hormone group of nuclear receptors and was shown to have a central role in prostate cancer development and progression. FOXA1, which was identified as downregulated in our analysis, is a pioneer TF that plays a pivotal role in partnering with AR to promote its attachment to chromatin. Remarkably, prior research revealed the role of FOXA1 as a suppressor of neuroendocrine differentiation and links its downregulation to the promotion of NEPC progression (31) and reprogrammed activity in NEPC (32). We further detected HNF4G as being significantly upregulated (adjusted p-value 0.0035, 1.77 log2 fold change) in the HAR cluster. This TF is generally involved in gastrointestinal neuroendocrine prostate cancer (GI-NEPC) and was shown to be expressed in 5% of primary prostate cancers and 30% of CRPCs. It is responsible for the activation of an AR-independent resistance mechanism involving the activation of gastrointestinal transcription and chromatin patterns (33). We also observed the presence of ARID1A and SMARCB1 among the top 5 differentially active TFs. Both of these TFs were shown to cooperate in the SWI/SNF chromatin remodeling complex and to be involved in prostate cancer lineage plasticity (34). Further, HOXB13, NKx3-1, FOXA2 and GATA2 are other TFs that are linked to NEPC (20,35,36) and were found to be differentially active. Lastly, our analysis highlighted TLE3 as a key player among the differentially active transcription factors whose loss was previously linked to the development of a glucocorticoid receptor (GR)-mediated resistance mechanism under androgen receptor inhibitors (37).

**Table 1:**
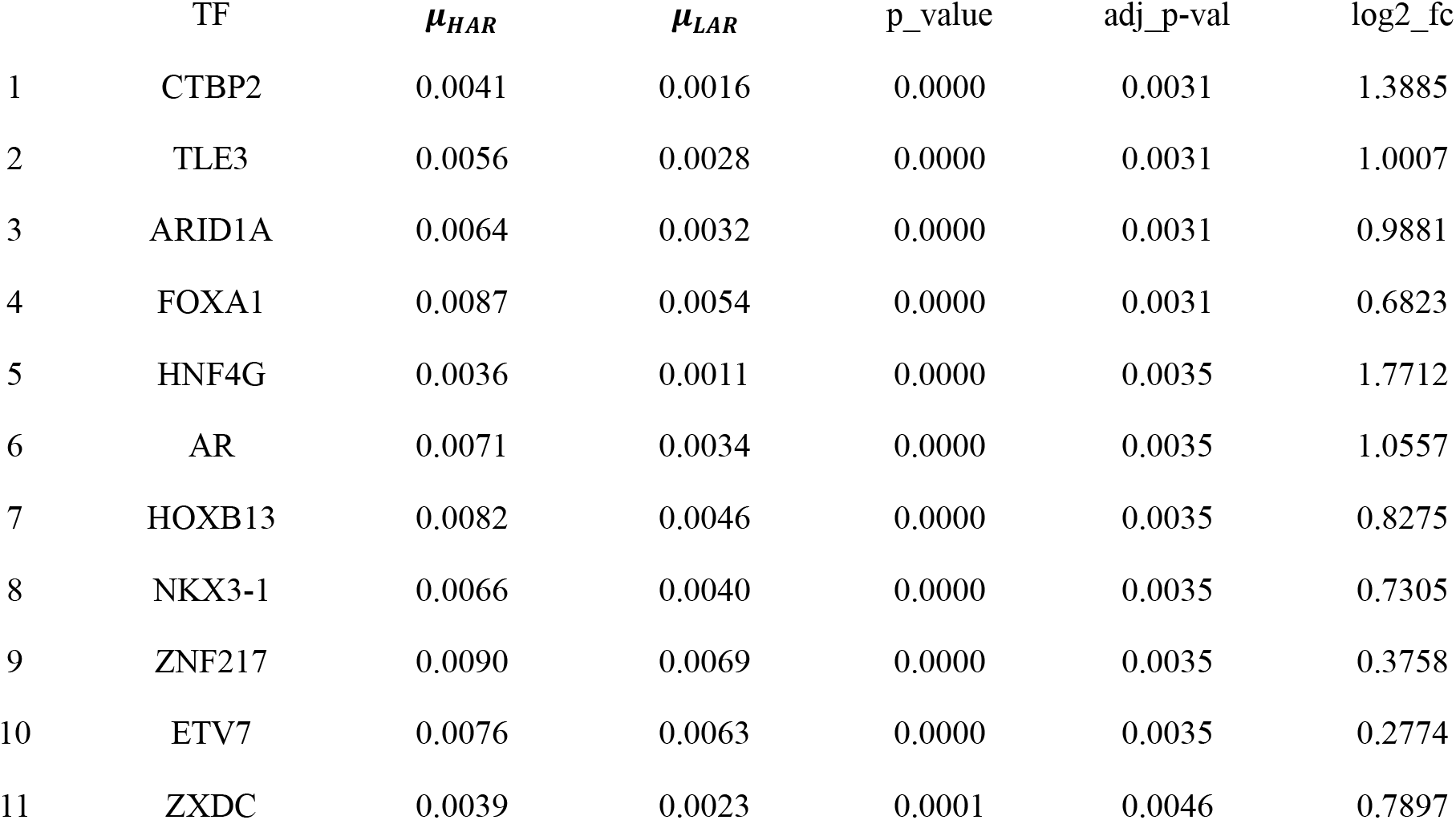
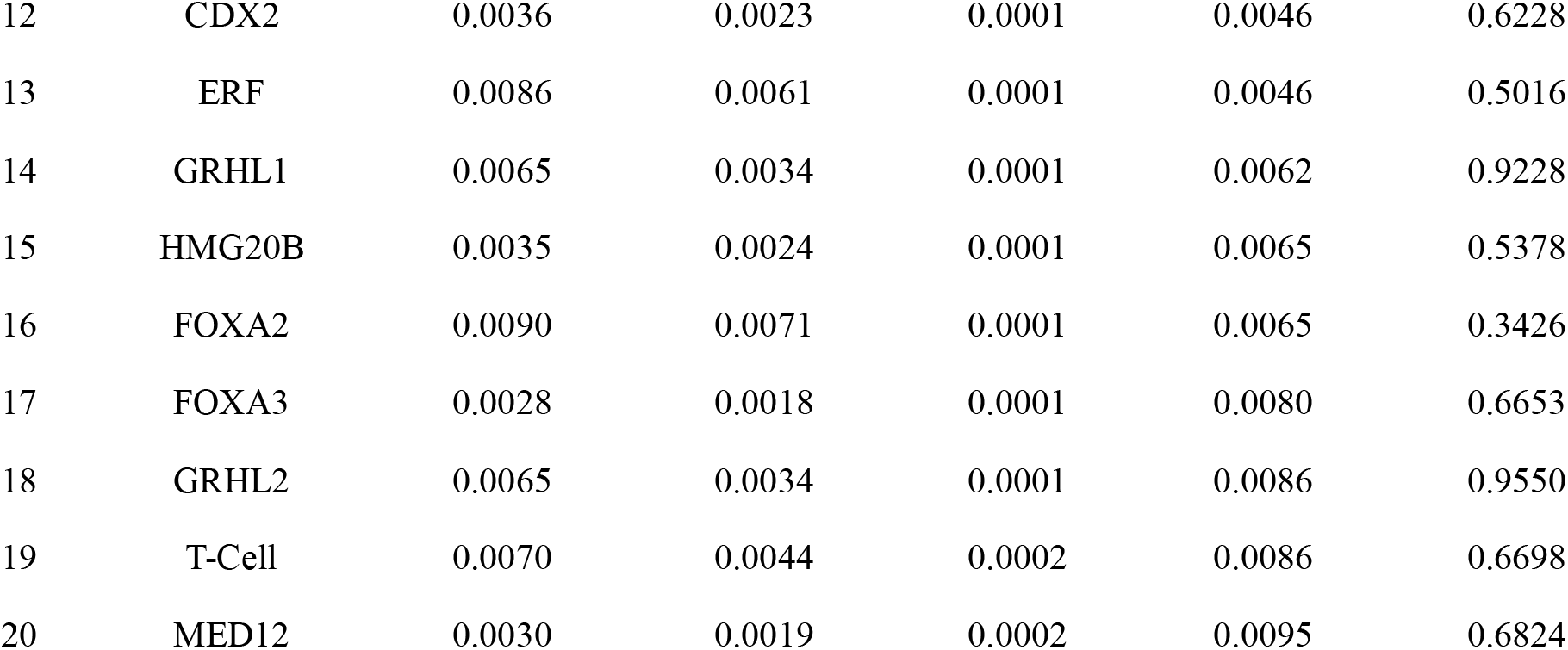
Results of the top 20 differentially active TF analysis performed on the HAR and LAR groups. (full table available in Supplementary Information). To include the direction of change, we calculated a pseudo log2 fold change, which retains the sign information.

### Enrichment analysis

After the analysis of differential activity, LBFextract performs an enrichment analysis step, which is carried out through the string API using all identified differentially active TFs. Through this step, the enrichment of the differentially active genes in different databases, which span from gene ontology to KEGG, REACTOME and WikiPathways, is retrieved. Here, we focused on the LAR and HAR clusters and found a significant enrichment in the “Androgen receptor signaling pathway”, alongside various processes associated with epithelial cell development and differentiation, prostate gland development, and neuron differentiation in the Gene Ontology process category. Moreover, the results of the DISEASES enrichment analysis contained many significant terms that are closely related to prostate cancer such as “prostate cancer”, “prostate carcinoma”, “prostate adenocarcinoma”, “adenocarcinoma”, and “reproductive organ disease”. The enrichment analysis in the TISSUES category indicated statistically significant enrichment in “whole blood” and “prostate epithelium cell line” as well as “prostate epithelium cell”. Furthermore, pathways including “Androgen receptor network in prostate cancer”, “Nuclear receptors” and “Endoderm differentiation” were found in the WikiPathways category. Additionally, pathways such as “Signaling by Nuclear Receptors,” “Estrogen-dependent gene expression”, “NR1H2 & NR1H3 regulate gene expression to limit cholesterol uptake”, and “NR1H2 & NR1H3 regulate gene expression linked to triglyceride lipolysis in adipose” in the REACTOME database were significantly enriched.

## DISCUSSION

We showcased the features and applications of our LBFextract package in a biomarker discovery setting for prostate cancer, highlighting its unique capabilities in feature extraction, differential activity analysis, and its plugin structure designed to meet the continuous growth in feature extraction methods in liquid biopsy research. In our package, we provide a default set of liquid biopsy feature extraction methods based on fragment coverage with major differences to coverage features extraction methods offered by other packages like SAMTools (38), Picard (39), DeepTools (40) Pysam (38,41,42), Mosdepth (43) and BEDTools (44,45).

While most of these general tools are based on read coverage or coverage in ranges, the coverage methods implemented in LBFextract are based on fragment coverage, in which the positions between read pairs are filled, and different strategies to increase the strength of the dyads’ derived signals are applied, making nucleosome phasing around active TFBSs more visible. (**Figure 2**b-d, Supp Figure 2). For example, we implemented midpoint coverage and middle-n points coverage (**Figure 2**c-d), previously used in (21) to increase the strength of the phasing signal, and central 60bp-coverage (8), which we reimplemented here without dependencies from other tools like fastx_trimmer, generalized to a user-defined central region, prevented double counting of overlapping reads and made it compatible with reads generated using less than 150 sequencing cycles, thus reducing sequencing requirements and costs.

Investigating the fragment length distribution signals derived after diverse *in silico* trimming strategies at CTCF TFBSs, an issue from using midpoint coverage with fragments having a length higher than 220bp becomes visible. Indeed, the center of dinuclesome-derived fragments is positioned between nucleosome-derived dyads. Unphased dinucleosomal and mononucleosomal fragments produce a weaker signal, which can be improved by better modelling the position of the dyad on polynucleosomal fragments. Therefore, we implemented the coverage around dyads signal, which provides a stronger dyad-derived signal. We modelled information about dyads coming from polynucleosomal structures, i.e. fragments having a length between multiples of the mono-nucleosomal length, thus increasing the amount of information used, which resulted in a stronger dyad-derived signal for phased nucleosomes (Supp Figure 2). With this strategy, we further improved upon the coverage analysis performed in our previous work (8). Indeed, for dinucleosome-derived fragments, an increasing bias concerning the location of the dyad is introduced the more a fragment becomes digested if only the central part of the read is considered, with the extreme case of fragments having less then 150bp, for which the positions are counted twice.

In our investigation, we examined the androgen receptor (AR) coverage signal within sample P148, which trans-differentiated into neuro-endocrine prostate cancer. Our results reveal a flat coverage profile at AR transcription factor binding sites (TFBSs), along with the disappearance of the recurrent peaks induced by nucleosome phasing in P148_3. These observations align with and support the conclusions drawn in prior studies (9,20) in which a similar behavior was described and shows the importance of liquid biopsy specific features. Indeed, with general coverage strategies, this information was not visible.

Finally, we expanded analyses that can be performed with LBFextract further into the potential fragmentomics space by implementing entropy derived features, which efficiently summarize variation in fragment length distribution signals. We used these features to infer the TF activity in PC and NEPC samples, showing that Relative Fragment Entropy (RFE) is powerful for recapitulating findings of our previous work (9,20). The validity of this signal is also supported by the fact that hematopoietic TFs like TAL1 and LYL1 show similar profiles between samples in contrast to condition specific TFs like AR, HOXB13 and FOXA1, which provide different signals in different conditions.

To identify potentially new TFs associated with these cancer subtypes, we extracted the coverage signal of 1058 TFs retrieved from the GTRD database, for all our high coverage prostate cancer samples. We performed cluster analysis and looked for differentially active TFs between the groups obtained with this approach, highlighting LBFextract’s capability of discovering both subtypes and subtype specific differentially active TFs from liquid biopsy data. Indeed, through this analysis we found a higher expression of the NKX3-1 TF in the HAR cluster which aligns with the finding that NKX3-1 is generally expressed in the luminal cell of the prostate where it is intertwined in regulative feed-forwards loop with the AR in both normal prostate and AR dependent prostate cancer (46), but may be downregulated or lost in neuroendocrine prostate cancer. We also observed a loss in the LAR group for the HOXB13, which is generally active in AR dependent PC and CRPC, but down-regulated in NEPC (47). This agrees with the low AR signal, low PSA values and high NSE values found in P148_3 and P198_5, which are part of the LAR cluster. Interestingly, two subunits of the mSWI/SNF complex, ARID1A and SMARCB1, were found to be downregulated in the LAR cluster suggesting a possible impairment of the BAF complex known to be involved in damage response. As suggested by Park Y. et al. (48), this can be exploited to challenge the tumor with PARP inhibitors combined with ionizing radiation.

## CONCLUSION

In this article, we introduced LBFextract, a Python package designed for the extraction and analysis of features from liquid biopsy data, with a specific emphasis on transcription factor-specific coverage and entropy features. A notable strength of LBFextract lies in its flexibility, allowing seamless integration of new feature extraction methods in the form of plugins, enabling adaptability to research approaches as needed. Our study demonstrated the capabilities of LBFextract in suggesting TFs for follow-up research and showcasing its ability to recapitulate signals observed in previous work. This validation was performed from both a coverage and fragmentomics perspective. We outlined the ability of identifying condition-specific TFs, demonstrating the tool’s utility in uncovering potential biomarkers in diverse biological contexts. LBFextract’s open architecture and compatibility with plugins seek to not only make it a standalone tool for reproducible feature extraction and biomarker identification, but also to contribute to the dynamic and collaborative nature of broader scientific and Python communities.

## ACKNOWLEDGEMENTS

We are especially grateful for the help and guidance of Prof. Michael Speicher, who passed away unexpectedly in September 2023. He was a great mentor, colleague, and friend. During his scientific career, he made many important contributions to the circulating cfDNA field, pioneering the analysis of WGS data of cfDNA and developing some of the earliest applications of these methods. His scientific passion will live on in our work. Special thanks to Hossein Hajiabolhassan, the other dedicated members of the BPDP group and Kircher lab, whose insights and collaboration have enriched the scientific depth of this work. We are very grateful to Prof. Ellen Heitzer for the critical review of this manuscript. We extend our sincere appreciation to Thomas Bauernhofer for generously providing the samples and clinical information used in this study. Figure 1 was created with BioRender.com.

