## Supplementary Information for "LBFextract: unveiling transcription factor dynamics from liquid biopsy data"

### Supplementary Methods

#### ***In silico* dilutions**

To *in silico* dilute the tumor samples, we created a pool of reads by merging 30 healthy male samples. Each sample was downsampled to 20x average coverage using *samtools*.

For each sample *I*, we counted the total number of reads, which we defined as *t*. We subsequently estimated the number of reads $r$ that needed to be sampled from *t* to achieve a specific tumor fraction, as shown in the following formula:

$$r_i = \left(\frac{s_d}{s_t}\right) t_i \qquad 22$$

in which $s_d$ represents the tumor fraction after dilution and $s_t$ the tumor fraction of the tumor sample. To infer $s_t$, ichorCNA (1) was used.

In this way, we were able to obtain the ratio required by *samtools view* to subsample the correct number of reads from the tumor sample. This was achieved using the following formula:

$$c_t = \frac{r_i}{t_i} \qquad 23$$

We calculated the number of reads to subsample from the pool of healthy reads $r_h$ and the ratio required by *samtools* to achieve this $c_h$in the following way:

$$r_h = t_i - r_i \qquad 24$$

$$c_h = \frac{r_h}{h} \qquad 25$$

In which *h* represents the number of reads in the healthy pool.

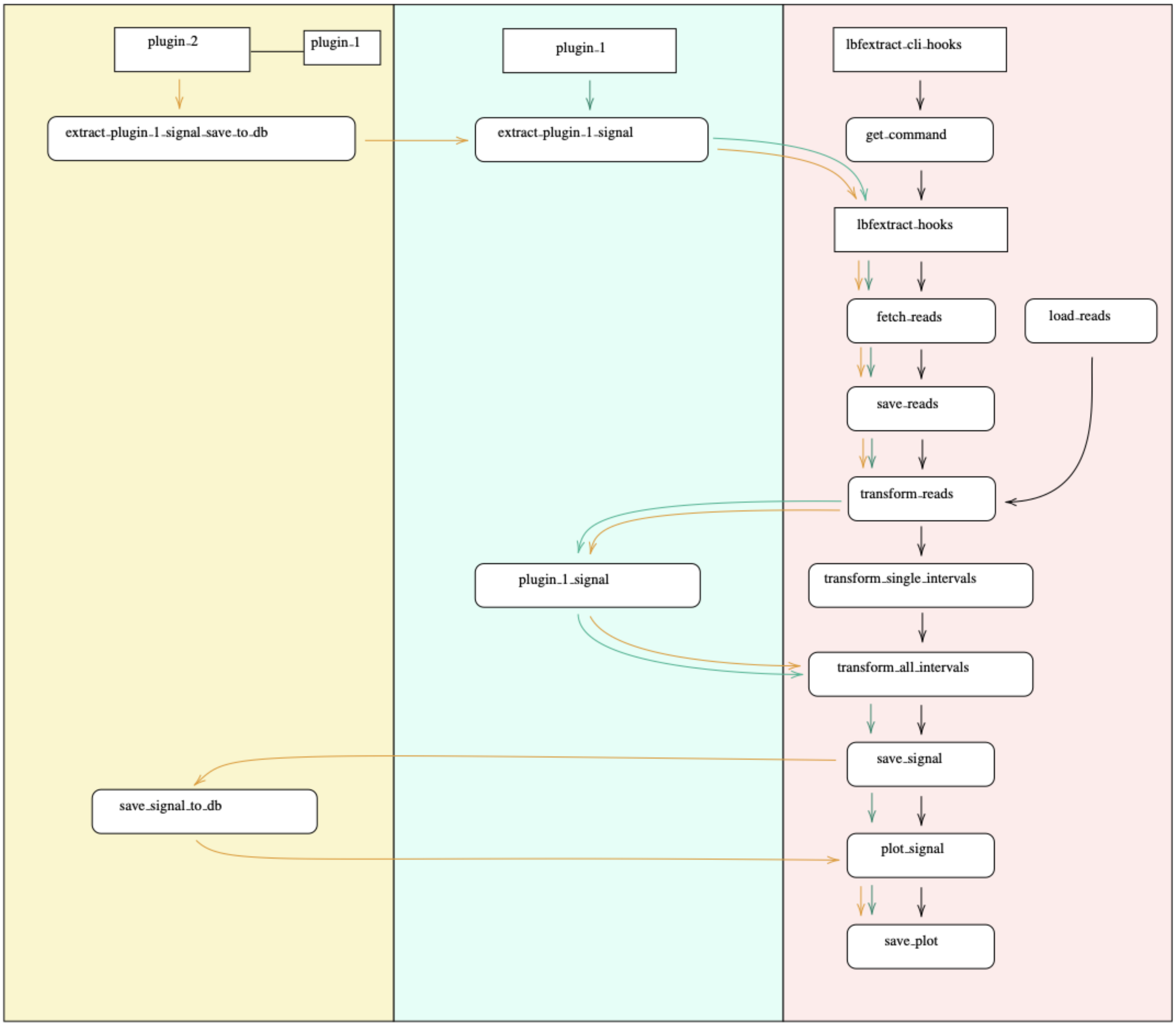


**Supp. Figure 1: Plugin interface of the LBFextract package.** The right column shows the workflow implemented by LBFextract with the hooks, which may be overwritten by other plugins to change LBFextract's behavior. The central column represents the hook implemented by "plugin_1" and the blue arrows represent the flow followed by the program. In this case "plugin_1" implements the transform_single_interval hook to extract a different signal and the flow follows the normal workflow till that point, at which it then enters the "plugin_1" hook "plugin_1_signal" returning to the original workflow afterword. The left column represents a second plugin, which extends the behavior of "plugin_1" to save the data to a database. In this case the program follows the workflow of "plugin_1" till the "save_signal" hook. Then, the program enters the "save_signal" hook implemented by "plugin_2".

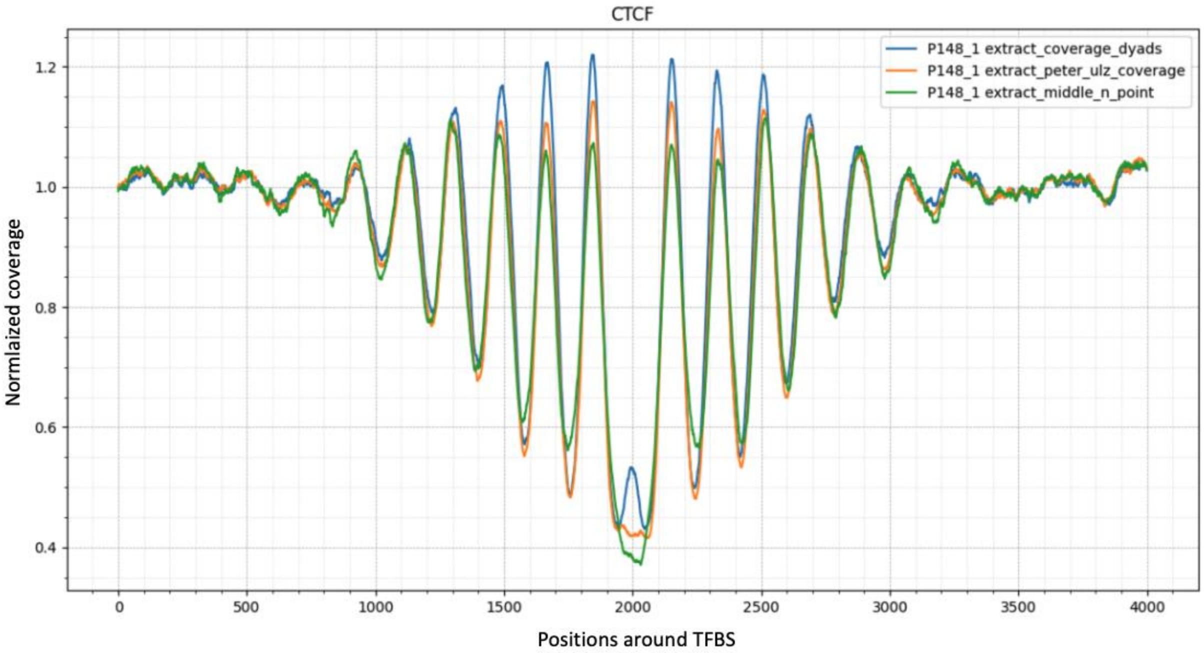


**Supp. Figure 2: Comparison of coverage around dyads, central 60bp-coverage and middle-n points coverage.** For middle-n points coverage and coverage around dyads a window of ± 30bp was used. For central 60bp-coverage, the default bases covering positions 53 to 113 from the start and the -53 to -113 from the end of each fragment were used.

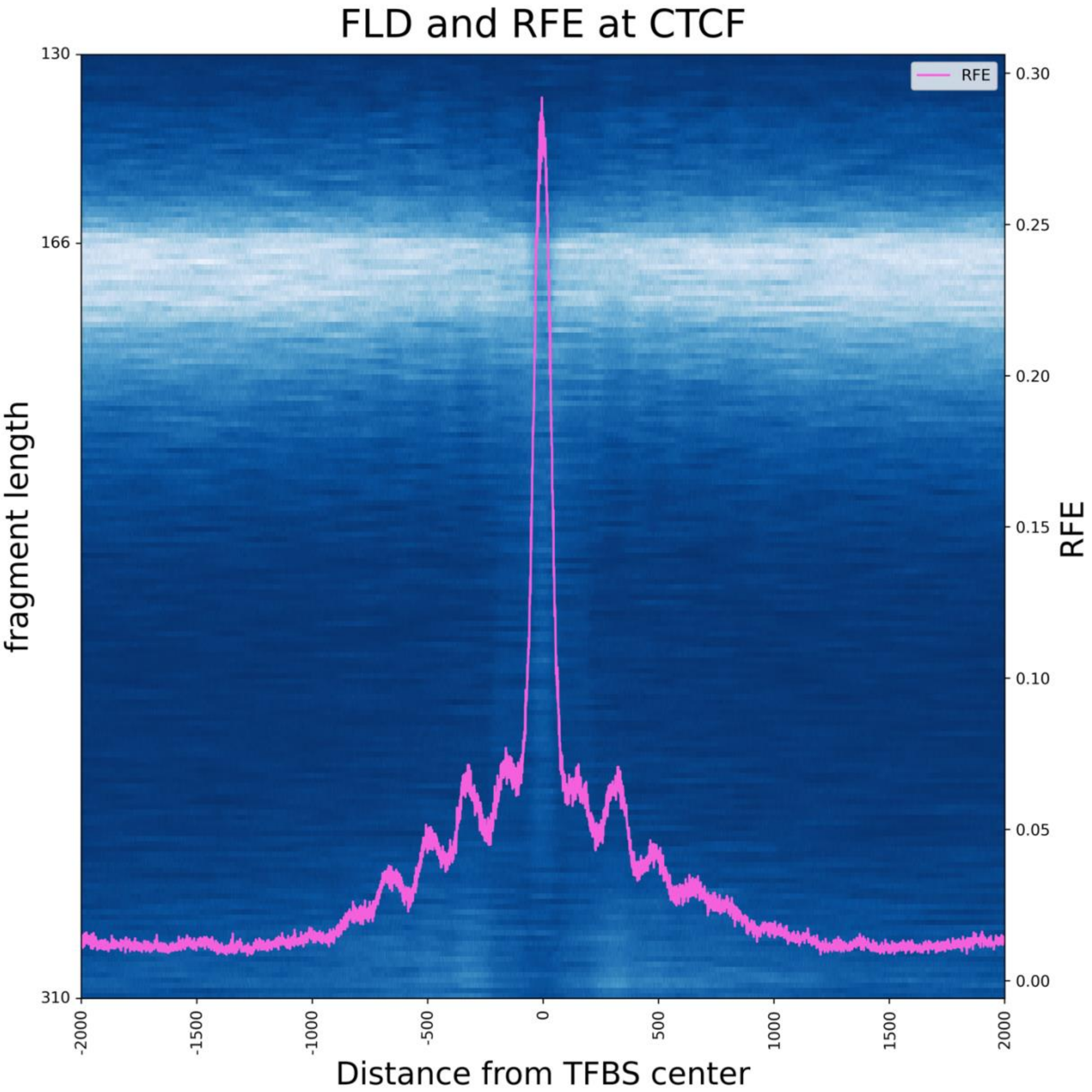


**Supp. Figure 3: relationship between RFE and FLD at CTCF.** RFE reaches its maximum where the fragment length distribution at a specific position is most different from the fragment length distribution in the flanking regions. In the case of TFBSs that have an open chromatin state like CTCF, this happens in the center.

**Clustering unsupervised and differentially active analyses**

Due to the heterogeneity of the HAR group, we also conducted k-means clustering to look for subclusters that may be present in this group. The selection of the best number of clusters was assessed using the elbow method (Supp. Figure 4), which identified three clusters. We calculated the differentially active TFs between these groups according to Algorithm 2.

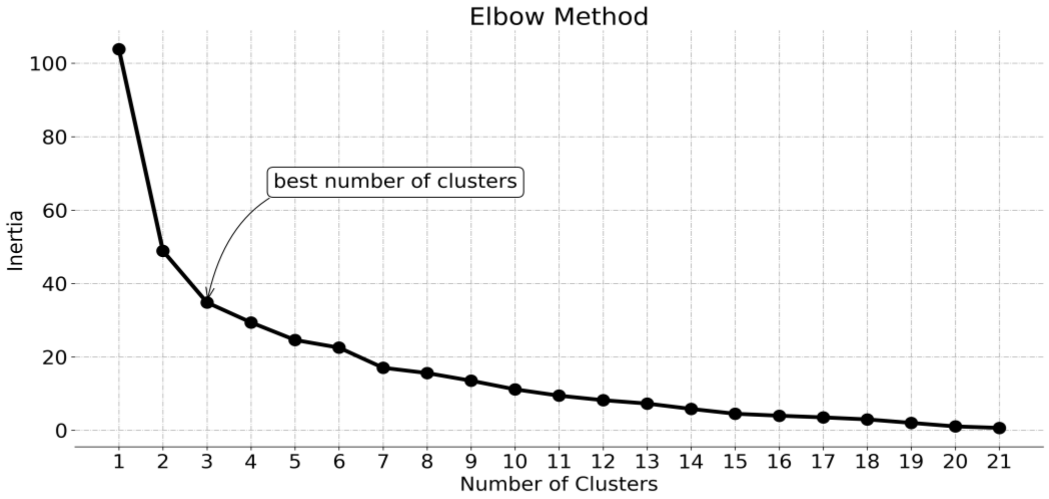


**Supp. Figure 4: Elbow method to select the number of clusters.** K-means was calculated varying the number of clusters from 1 to 21. With each number of clusters, we measured the inertia and selected the best number of clusters as the point at which the curve starts to flatten.

Notably, clusters IA and HA, which were found with k-means clustering, exhibited an overlap with the HAR cluster (Figure 5), while cluster LA overlapped with the LAR group (Figure 5). This characteristic was also reflected in the differentially active analysis between these groups. Specifically, the differentially active TFs identified between HA - IA and IA - LA were subsets of those identified in the LA - HA comparison. We therefore looked at the difference in differentially active TFs found in the comparisons of groups HA - IA and IA - LA and observed the presence of many TFs part of the AR network, such as ELK1, STAT3, ETV4, SP1, BRCA1 as well as TFs involved in the regulation of fatty acids, such as NR1H3 and NR1H2, which are only present in the list of differentially active TFs obtained by the comparison between the LA and HA groups.

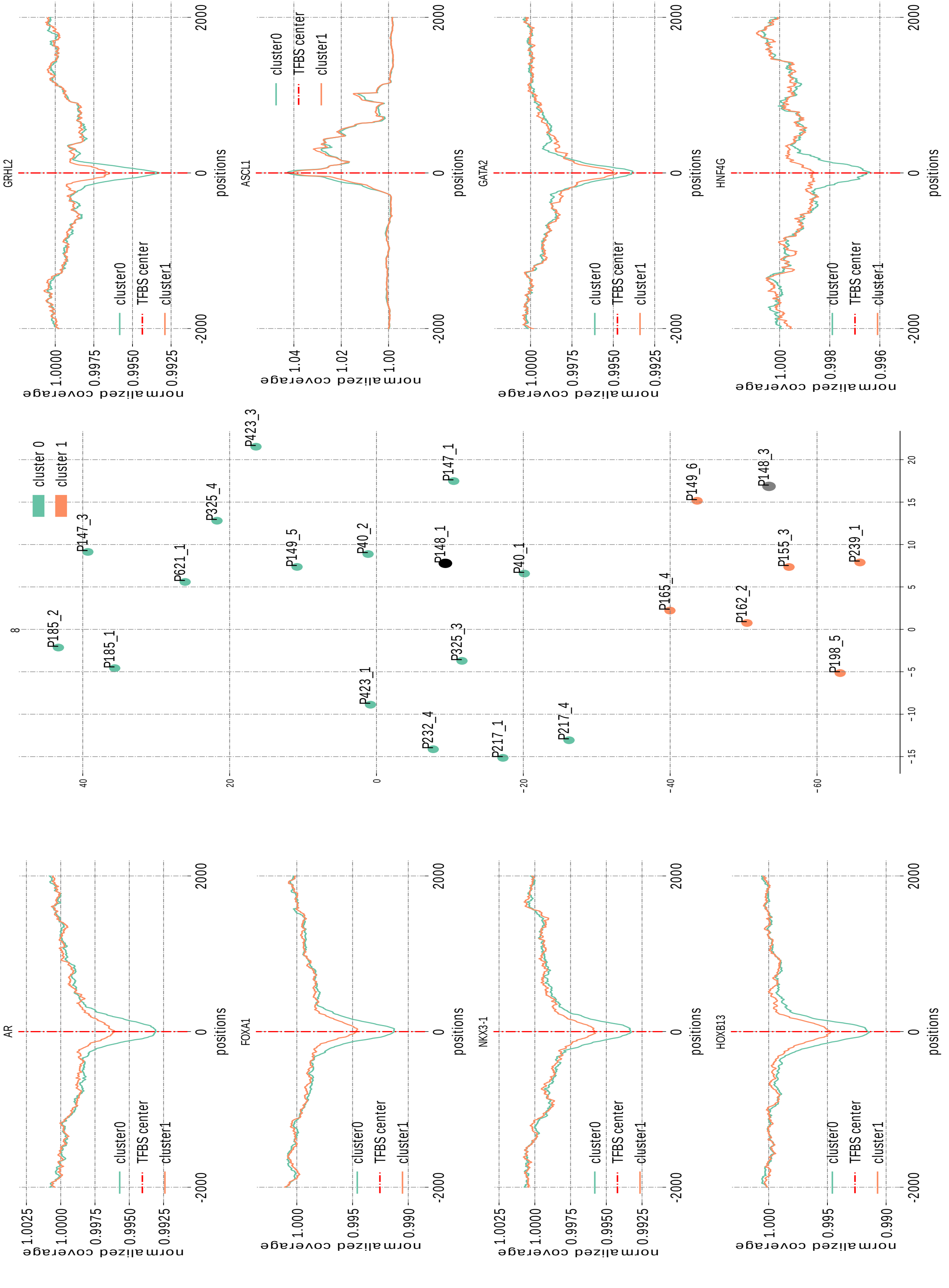


**Supp. Figure 5: Coverage signal of the selected TFBSs used for clustering and distribution of the sample-specific signals in the t-SNE-embedded latent space.** In the left and right columns, coverage signals used in the semi-supervised clustering of the TFs are visualized for the two identified clusters. In the central column, we visualized the samples using a t-SNE projection

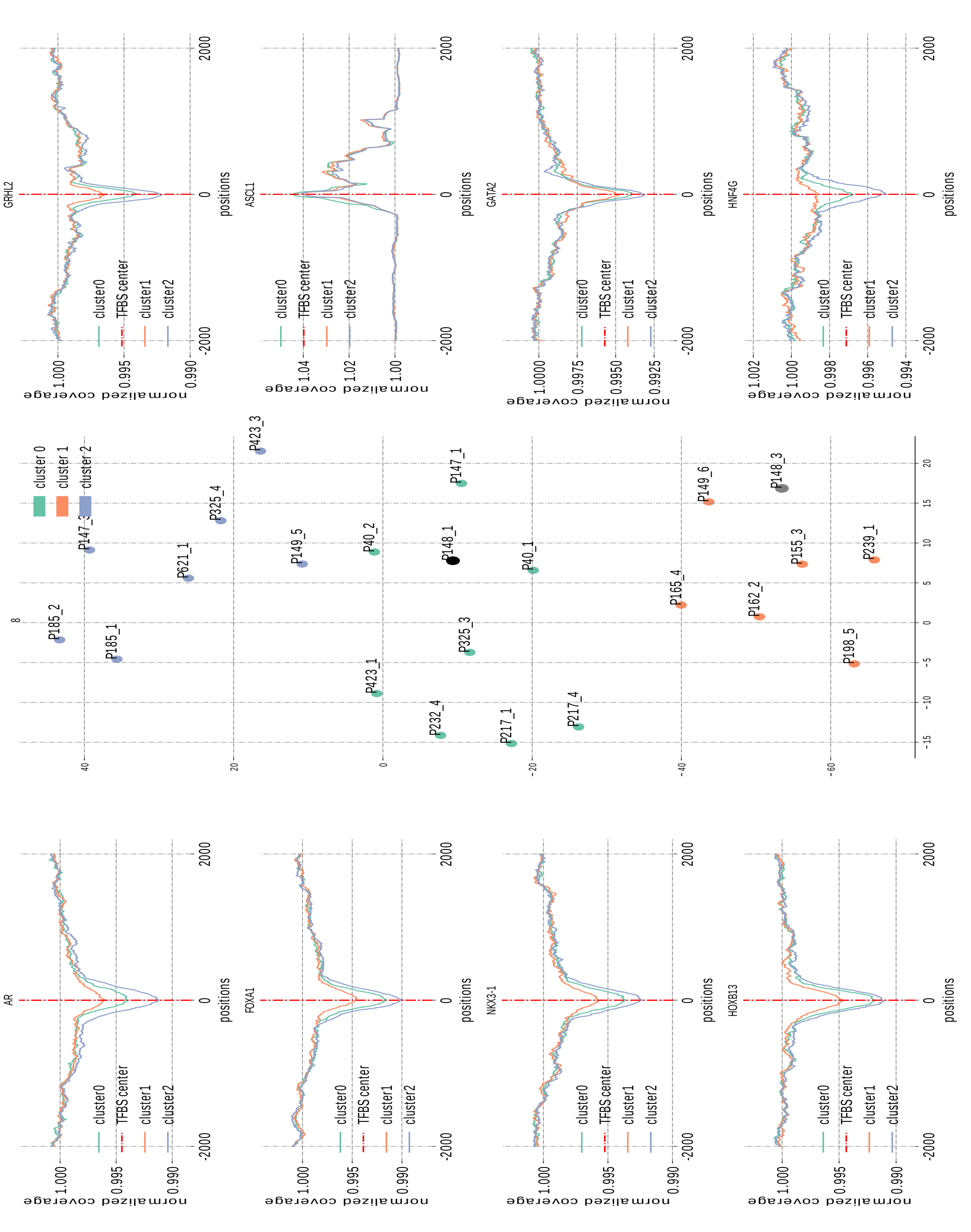


**Supp. Figure 6: Coverage signal of the selected TFBSs used for clustering and distribution of the sample-specific signals in the t-SNE-embedded latent space.** In the left and right columns, coverage signals used in k-means clustering are visualized for the three identified clusters. In the central column, we visualized the samples using a t-SNE projection.

| | TF | $\mu_{HAR}$ | $\mu_{LAR}$ | p_value | adj_p-val | log2_fc |
|---|---|---|---|---|---|---|
| 1 | CTBP2 | 0.0041 | 0.0016 | 0.0000 | 0.0031 | 1.3885 |
| 2 | TLE3 | 0.0056 | 0.0028 | 0.0000 | 0.0031 | 1.0007 |
| 3 | ARID1A | 0.0064 | 0.0032 | 0.0000 | 0.0031 | 0.9881 |
| 4 | FOXA1 | 0.0087 | 0.0054 | 0.0000 | 0.0031 | 0.6823 |
| 5 | HNF4G | 0.0036 | 0.0011 | 0.0000 | 0.0035 | 1.7712 |
| 6 | AR | 0.0071 | 0.0034 | 0.0000 | 0.0035 | 1.0557 |
| 7 | HOXB13 | 0.0082 | 0.0046 | 0.0000 | 0.0035 | 0.8275 |
| 8 | NKX3-1 | 0.0066 | 0.0040 | 0.0000 | 0.0035 | 0.7305 |
| 9 | ZNF217 | 0.0090 | 0.0069 | 0.0000 | 0.0035 | 0.3758 |
| 10 | ETV7 | 0.0076 | 0.0063 | 0.0000 | 0.0035 | 0.2774 |
| 11 | ZXDC | 0.0039 | 0.0023 | 0.0001 | 0.0046 | 0.7897 |
| 12 | CDX2 | 0.0036 | 0.0023 | 0.0001 | 0.0046 | 0.6228 |
| 13 | ERF | 0.0086 | 0.0061 | 0.0001 | 0.0046 | 0.5016 |
| 14 | GRHL1 | 0.0065 | 0.0034 | 0.0001 | 0.0062 | 0.9228 |
| 15 | HMG20B | 0.0035 | 0.0024 | 0.0001 | 0.0065 | 0.5378 |
| 16 | FOXA2 | 0.0090 | 0.0071 | 0.0001 | 0.0065 | 0.3426 |
| 17 | FOXA3 | 0.0028 | 0.0018 | 0.0001 | 0.0080 | 0.6653 |
| 18 | GRHL2 | 0.0065 | 0.0034 | 0.0001 | 0.0086 | 0.9550 |
| 19 | T-Cell | 0.0070 | 0.0044 | 0.0002 | 0.0086 | 0.6698 |
| 20 | MED12 | 0.0030 | 0.0019 | 0.0002 | 0.0095 | 0.6824 |
| 21 | ZFAT | 0.0041 | 0.0026 | 0.0002 | 0.0095 | 0.6614 |
| 22 | KMT2C | 0.0122 | 0.0105 | 0.0002 | 0.0095 | 0.2176 |
| 23 | THAP1 | 0.0203 | 0.0186 | 0.0003 | 0.0158 | 0.1273 |
| 24 | SMARCB1 | 0.0061 | 0.0039 | 0.0004 | 0.0160 | 0.6357 |
| 25 | ETV2 | 0.0055 | 0.0043 | 0.0004 | 0.0160 | 0.3569 |
| 26 | SAP130 | 0.0048 | 0.0040 | 0.0004 | 0.0160 | 0.2407 |
| 27 | ETV4 | 0.0096 | 0.0085 | 0.0005 | 0.0200 | 0.1686 |
| 28 | DLX4 | 0.0066 | 0.0053 | 0.0005 | 0.0203 | 0.3156 |

| | | | | | | |
|---|---|---|---|---|---|---|
| 29 | AHR | 0.0017 | 0.0004 | 0.0007 | 0.0235 | 2.2838 |
| 30 | TBP | 0.0185 | 0.0165 | 0.0007 | 0.0235 | 0.1594 |
| 31 | DEAF1 | 0.0163 | 0.0147 | 0.0007 | 0.0235 | 0.1521 |
| 32 | KLF3 | 0.0033 | 0.0020 | 0.0008 | 0.0241 | 0.6688 |
| 33 | DLX2 | 0.0046 | 0.0029 | 0.0008 | 0.0241 | 0.6551 |
| 34 | RBM25 | 0.0049 | 0.0056 | 0.0008 | 0.0241 | -0.2088 |
| 35 | NR1H3 | 0.0026 | 0.0013 | 0.0011 | 0.0308 | 1.0090 |
| 36 | ZNF843 | 0.0024 | 0.0017 | 0.0011 | 0.0308 | 0.5348 |
| 37 | GATA2 | 0.0061 | 0.0047 | 0.0011 | 0.0308 | 0.3835 |
| 38 | GTF2B | 0.0135 | 0.0124 | 0.0011 | 0.0308 | 0.1186 |
| 39 | MIER3 | 0.0077 | 0.0068 | 0.0012 | 0.0320 | 0.1940 |
| 40 | FXR2 | 0.0019 | 0.0011 | 0.0015 | 0.0381 | 0.7077 |
| 41 | ETV1 | 0.0209 | 0.0191 | 0.0015 | 0.0381 | 0.1312 |
| 42 | USF2 | 0.0209 | 0.0193 | 0.0015 | 0.0381 | 0.1183 |
| 43 | GREB1 | 0.0012 | -0.0006 | 0.0015 | 0.0381 | -0.9677 |
| 44 | CUL4A | -0.0045 | -0.0031 | 0.0016 | 0.0381 | 0.5532 |
| 45 | L3MBTL4 | 0.0057 | 0.0049 | 0.0016 | 0.0381 | 0.2312 |
| 46 | ZNF511 | 0.0098 | 0.0085 | 0.0017 | 0.0386 | 0.1960 |
| 47 | KAT8 | 0.0100 | 0.0090 | 0.0017 | 0.0386 | 0.1547 |
| 48 | KLF5 | 0.0174 | 0.0155 | 0.0019 | 0.0403 | 0.1666 |
| 49 | PBX3 | 0.0219 | 0.0201 | 0.0019 | 0.0403 | 0.1224 |
| 50 | OVOL3 | 0.0042 | 0.0034 | 0.0020 | 0.0416 | 0.3159 |
| 51 | EHF | 0.0124 | 0.0110 | 0.0020 | 0.0416 | 0.1770 |
| 52 | NR1H2 | 0.0064 | 0.0051 | 0.0024 | 0.0475 | 0.3129 |
| 53 | TFE3 | 0.0048 | 0.0040 | 0.0024 | 0.0475 | 0.2688 |
| 54 | MEIS3P1 | -0.0127 | -0.0202 | 0.0024 | 0.0475 | -0.6765 |

**Table 1: Results of the top 20 differentially active TF analysis performed on the HAR and LAR groups.** To include the direction of change, we calculated a pseudo log2 fold change, which retains the sign information.

**Algorithm 1:** Coverage around dyads

**Data:** List $F$ of fragments
**Data:** Window $w$ around dyad center
**Result:** coverage $C$

1 $S \leftarrow []$ ;
2 $d \leftarrow get\_fld$(chr12:34300000-34500000);
3 $p \leftarrow get\_peak(d)$ ;
4 **for** *each fragment* $f$ *in* $F$ **do**
5     $n \leftarrow \left\lfloor \frac{|f|}{p} \right\rfloor$ ;
6     $r \leftarrow f \bmod p$;
7     $p_n \leftarrow (|f| - r)|f|^{-1}$;
8     $p_{n+1} \leftarrow 1 - p_n$;
9     **if** $p_n > p_{n+1}$ **then**
10         $v \leftarrow n$;
11     **else**
12         $v \leftarrow n+1$;
13     $f_e \leftarrow v * p$;
14     $f_m \leftarrow \left\lfloor \frac{|f|}{2} \right\rfloor$;
15     $f_s \leftarrow f_m - \left\lfloor \frac{|f_m|}{2} \right\rfloor$;
16     $f_e \leftarrow f_m + \left\lfloor \frac{|f_m|}{2} \right\rfloor$;
17     $f \leftarrow [f_s, f_e)$ ;
18     $f_{dyad} \leftarrow []$;
19     $m \leftarrow \left\lfloor \frac{|f|}{(v*2)} \right\rfloor$;
20     **for** $i \leftarrow 1$ **to** $v*2$ **by** $2$ **do**
21         $f_{dyad}$.append($f_{[f_s+(m*i)-w, f_s+(m*i)+w)}$) ;
22     S.append($f_{dyad}$)
23 $C = coverage(S)$;
24 **return** $C$;

**Algorithm 1: Algorithm to extract the coverage around dyads from a set of fragments.** We model the probability of each fragment coming from a n or $n+1$ polynucleosomal structure (lines 7-8). This is used to reconstruct the size of each fragment before degradation (lines 13-17), which in turn is used to better place the position of the dyad and a stronger nucleosome derived signal (lines 18-22).

**Algorithm 2:** From bam files to differentially active TFs

**Input:** $\mathbb{B}=\{bam\_file_i : i \in [1,\#samples]\}$
**Input:** $f$ : set_of_bam_files $\rightarrow \mathbb{Y}^{n\times m}$
**Input:** $G=\{i : i \in [1,\#groups]\}$
**Input:** $\mathbf{l}=(i_1,i_2,i_3,\ldots,i_n), i \in G$
**Input:** $n$ = number_of_transcription_factors
**Input:** $m$ = number_of_positions
**Input:** $F=\{i : i \in [1,\#transcription\_factors]\}$

1 **Function** `GetDiffActiveTf`($\mathbb{B}, f, \mathbf{l}, n, m, F$)**:**
2 $\quad S=\mathbf{0}^{n\times n\_groups}$ ;
3 $\quad groups=\{i : i \in l\}$ ;
4 $\quad n\_groups=\#groups$ ;
5 $\quad a=\{\mathbf{0}_i^n : i \in [1,\#F]\}$ ;
6 $\quad T \leftarrow \{0_i^{n\times m\times t} : i \in [1,\#samples]\}$;
7 $\quad$ **for** $i$ *in* $[1,\#samples]$ **do**
8 $\quad\quad$ **for** *t in F* **do**
9 $\quad\quad\quad T_{[i,:,t]} \leftarrow f_t(get\_regions\_from\_bam(\mathbb{B}_i))$ ;
10 $\quad$ **if** $n\_groups > 2$ **then**
11 $\quad\quad$ **for** $t$ *in* $F$ **do**
12 $\quad\quad\quad a[t] \leftarrow get\_accessibilities\_per\_group(T_{[:,:,t]})$ ;
13 $\quad\quad\quad P_{kruskal}[t] \leftarrow Kruskal\_Wallis(a[t],\mathbf{l})$ ;
14 $\quad\quad P_{corrected\_kruskal} \leftarrow MultipleTestCorrection(P_{kruskal})$ ;
15 $\quad\quad$ **for** $t$ *in* $F$ **do**
16 $\quad\quad\quad$ **if** $P_{kruskal} < \alpha$ **then**
17 $\quad\quad\quad\quad P_{dunn} \leftarrow posthoc\_dunn(a[t])$ ;
18 $\quad\quad\quad P_{corrected} \leftarrow MultipleTestCorrection(P_{dunn})$;
19 $\quad$ **else**
20 $\quad\quad$ **for** $t$ *in* $F$ **do**
21 $\quad\quad\quad a[t] \leftarrow get\_accessibilities\_per\_group(T_{[:,:,t]})$ ;
22 $\quad\quad\quad P_{mwu}[t] \leftarrow Mann\_Whitney\_U\_test(a[t],\mathbf{l})$ ;
23 $\quad\quad P_{corrected} \leftarrow MultipleTestCorrection(P_{mwu})$ ;
24 $\quad S \leftarrow P_{corrected}$ ;
25 $\quad$ **return** $S$ ;

**Algorithm 2: Algorithm to extract differentially active transcription factors from signals at transcription factor binding sites (TFBSs).** We extract the features f for each transcription factor *t* from the BAM file of sample *i* (lines 7-9). Subsequently, we calculate the accessibility of each feature and, in case of more than 2 groups, apply the Kruskal Wallis test (lines 10-18), Mann

Whitney U Test (line 20-23) otherwise. Because of multiple groups and multiple TFs to be considered, multiple tests correction is applied.
